# The role of cellular filamentation in bacterial aggregation and cluster-cluster assembly

**DOI:** 10.1101/2023.09.01.555911

**Authors:** Samuel Charlton, Gavin Melaugh, Davide Marenduzzo, Cait MacPhee, Eleonora Secchi

## Abstract

Bacterial aggregate formation and surface accumulation are increasingly viewed as alternative pathways for biofilm colonization. However, little is known about the dynamics of bacterial aggregate cluster-cluster assembly and their subsequent microstructural and mechanical properties. To this end, we studied experimentally and computationally an aggregating bacterial system that forms a space-spanning interconnected network via cluster-cluster assembly. By controllably inducing bacterial filamentation, we aimed to understand how cell length distribution and cell surface hydrophobicity control the dynamics of aggregation and sedimentation, as well as the microstructure and mechanics of the settled bacterial networks. We found that filamentation lowers the percolation threshold, leading to gelation at a lower number density with distinct assembly dynamics and lower network connectivity. Furthermore, we analyzed the mechanical properties of the bacterial networks. Static stress tests reveal three yielding modes: discrete cluster-cluster disassembly, collective delamination, and sub-regional network fracture. The yielding modes are consistent with the gel-like viscoelastic properties of the cluster-cluster assembled networks observed during macroscale rheometry. In particular, we observe a scaling relationship between the storage modulus and the volume fraction, characteristic of an attractive rod gel. Our experimental observations are supported by Langevin dynamic simulations, providing mechanistic insights into the factors determining network self-assembly and connectivity. Our findings elucidate the gel-like structure-function dynamics in cluster-cluster aggregated bacterial systems and underscore the fundamental importance of filamentation in their properties and mechanical behavior.

## Introduction

Bacteria can assemble to form collective structures, either as surface-attached biofilms or suspended non-surface associated aggregates (*1*). The formation and subsequent gravitational settling of bacterial aggregates via encounters or growth is increasingly recognised as an alternative pathway to biofilm formation (*2–5*). This process, known as bioflocculation, is widely utilized in biotechnology applications, such as bioreactor operation and water remediation (*6*). Yet, how bacterial physiology, e.g. cell length, tunes the formation dynamics of bacterial aggregates and the structure-function relationships of their subsequent surface-settled structures remains underexplored.

It is known that bacteria can regulate their cell length through a process known as conditional filamentation. This process is triggered by various environmental stressors, for example, nutrient depletion and antibiotic exposure (*7–9*). Filamentation offers several biophysical advantages in stressed environments, including better access to nutrients in patchy environments and enhanced colonisation of nutrient-rich particles (*10–13*). Cell length geometrically influences encounter rates, which, together with cell-cell cohesion strength, control bioflocculation (*14*); however, the impact of filamentation on aggregation and their subsequent network formation has not yet been systematically investigated across different length scales within the same system.

In abiotic monodisperse colloidal systems, the aspect ratio of the particles and the system interaction strength are fundamental parameters that determine aggregation and the formation of space spanning colloidal gel networks (*15, 16*). In these systems, cluster-cluster aggregation is a prerequisite for network formation (*17*). Here, the aspect ratio controls the encounter rate between particles (*18*), while the strength of interparticle interaction, which confers mechanical stability to gelated networks, can be modulated through mechanisms such as depletion attraction, polymer bridging, and hydrophobic interactions (*19, 20*). Colloidal networks display various emergent properties, including viscoelastic rheology and time-dependent collapse, in which the strength of the network is overcome by gravitational settling (*21, 22*). Although these structural and dynamical principles are well understood in monodisperse nonliving systems, they have yet to be robustly investigated in filamenting aggregating bacterial systems, which are inherently polydisperse in cell length.

Various assembly mechanisms observed in colloidal systems have been identified as playing a role in the assembly of bacterial communities. For example, bacterial aggregation in polysaccharide-secreting cells can occur through entropic depletion attraction, leading to phase separation and spatial heterogeneity in biofilm colonies (*23, 24*). In the gut microbiota, the size dependence of bacterial aggregates can be modeled by combining colloidal gel theory with bacterial growth dynamics (*25*). The viscoelastic structure-function relationships consistently found in biofilms exhibit behaviors analogous to those of colloidal polymer gel composites (*26–28*). Finally, in the case of biofilm morphogenesis from single cells and in response to mechanical shear, biofilm architecture can be understood using interaction potentials and mechanical contact models (*29–32*).

Here, we present a multi-scale study quantifying the impact of cellular filamentation on the aggregation dynamics and hierarchical cluster-cluster assembly in a bacterial system exhibiting bioflocculation. Specifically, we use *Comamonas denitrificans*, a bacterium capable of inducible, conditional filamentation. Using the dissolved oxygen concentration during growth as a lever to control the distribution of cell lengths, we elucidate the influence of cell length on the assembled bacterial networks, quantifying both the microstructure and mechanical behaviour. By performing Langevin dynamics simulations, we map the structural and dynamical properties observed in the networks formed in our simulations to those in our experimental system. Our findings provide new insights into the assembly of surface-associated biofilms through cluster-cluster aggregation, highlighting the biophysical mechanisms that shape their network properties and modulate their mechanical behaviour.

## Results

### Oxygen depletion induces controllable filamentation

To systematically quantify the impact of cell length on aggregation and subsequent surface-associated cluster-cluster assembly, we select a bacterium displaying both filamentation and aggregation. *C. denitrificans* is a biofilm-forming, gram-negative bacterium known to seamlessly switch between aerobic and anoxic growth conditions (*33*). This bacterium plays an important role in bioaugmentation and bioremediation (*34, 35*), and exhibits conditional filamentation.

Filamentation is known to be induced by several environmental factors (*7*). Therefore to screen for levers that induce filamentation in *C. denitrificans* in the liquid culture we quantify how dissolved oxygen (DO) concentration, incubation temperature, and the presence or absence of shaking impact cell length distribution after 24 hours incubation (see Methods). Regardless of whether the culture is grown in stationary or shaking conditions, *C. denitrificans* exhibits a propensity toward filamentation under oxygen-limited conditions, corresponding to a reduced headspace (Fig. 1a). Filamentation is also induced by adjusting the culture temperature, particularly in oxygen-limited conditions (Figs. 1b and c).

**Figure 1:**
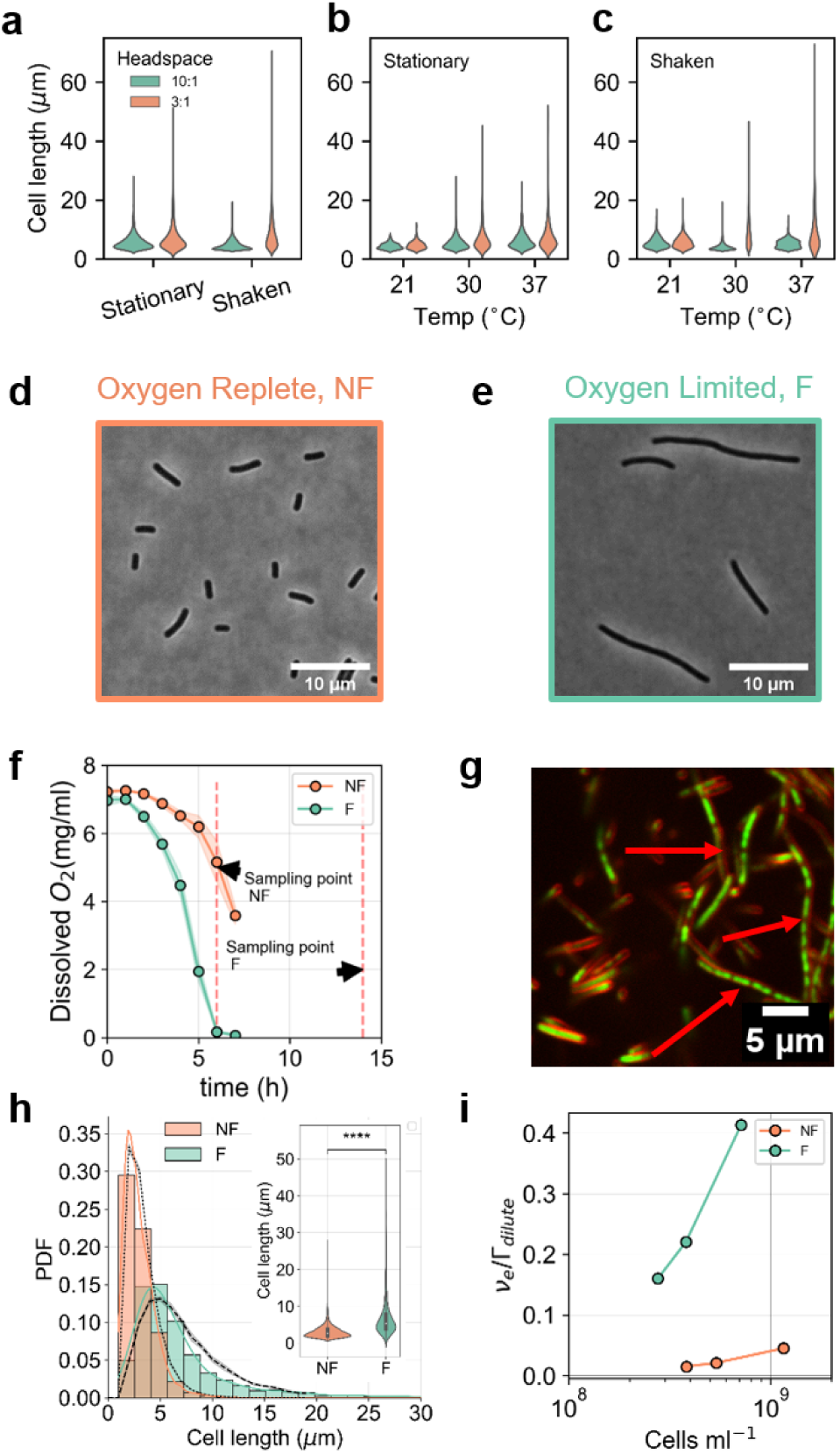
Filamentation can be tuned by controlling oxygen concentration. (**a-c**) Cumulative cell length distribution at low (3:1) and high (10:1) headspaces in stationary and shaken culture conditions after 24 hours incubation at 30^◦^C (a), after 24 hours incubated at low (21^◦^C), optimal growth (30^◦^C) and high incubation temperature (37^◦^C) in stationary (b) and shaken conditions (c). (**d, e**) Representative phase contrast images of *C. denitrificans* grown in oxygenreplete and oxygen-limited conditions, respectively, and sampled at the points annotated in (**f**). (g) Dissolved O_2_ concentrations in the F and NF culture vessels during cell growth. The red dashed lines and arrows indicate the sampling point for the cells where cell densities were equal. Presented are the mean of 3 biological replicates plus minus the standard deviation. (h) Representative CLSM image of F *C. denitrificans*. The red arrows highlight the multinucleated filamentous structure, whereby multiple cell cytoplasms (greenlabeled with Syto 9) are encased within a continuous cell membrane (red labeled with SynaptoRed C2). (**h**) Cell length probability distribution functions (PDF) for the NF (n = 10117) and F (n = 3570) cell suspensions. Cell lengths were calculated fromphase contrast microscopy images like the ones

Based upon our parameter screen, filamentation is most strongly induced by controlling oxygenation conditions during culture (Figs. 1a-c, see Methods). Therefore, to produce cell suspensions with repeatable, controllable cell length distributions we maintain a constant incubation temperature of 30^◦^ C and vary the dissolved oxygen concentration. In the filamentous (F) phenotype (Fig. 1d, e), the cells are significantly longer and exhibit a greater polydispersity in cell length (the standard deviation over the mean changes from 1.27 vs 0.34) compared to the non-filamentous (NF) phenotype (Fig. 1h inset). The cell length distributions for the NF and F phenotypes follow a logarithmic distribution, indicative of cumulative fragmented division events that occur during cell growth (Fig. 1h and (*36*)). Modulation of oxygenation has a negligible impact on cell viability, as confirmed by live-dead assay (fig. S1), and the filaments contain multiple nucleoids, as observed in confocal laser scanning microscopy (CLSM) images (Fig. 1g).

In monodisperse suspensions of rod-shaped particles or polymers, number density is typically used as a metric relating packing and entanglement (*37, 38*). However, in polydisperse cellular suspensions, the length distribution must also be considered (Fig. 1h). Therefore, we compute the effective number density, *ν_e_* (see Methods). We normalise *ν_e_* by the respective upper boundaries of the theoretical dilute regime Γ*_dilute_*, where bacteria begin to interact sterically as their gyration radii overlap. Our results show that filamentation leads to a 20-fold increase in the effective number density at comparable cell concentrations (Fig. 1i).

### Cell suspensions of *C. denitrificans* aggregate and display gel-like dynamics upon sedimentation

We first investigate how the induction of filamentation influences the dynamics of macroscale aggregation and settling in our bacterial system. For these experiments, we place F and NF suspensions in macroscopic optical cuvettes (17 mm × 5 mm × 10 mm) and image them through time using a horizontal imaging platform (see Methods). Regardless of phenotype, we observe that stationary phase suspensions of *C. denitrificans* aggregate and form gel-like networks via cluster-cluster aggregation, which sediment under gravity. Assembly occurs in three distinct stages: 1) an initial quiescent phase of duration *t_q_*; 2) phase separation; and 3) compaction (Fig. 2a, b). To quantify the system dynamics, we track the position of the interface that emerges between the concentrated and supernatant phases (Fig. 2a, b).

**Figure 2:**
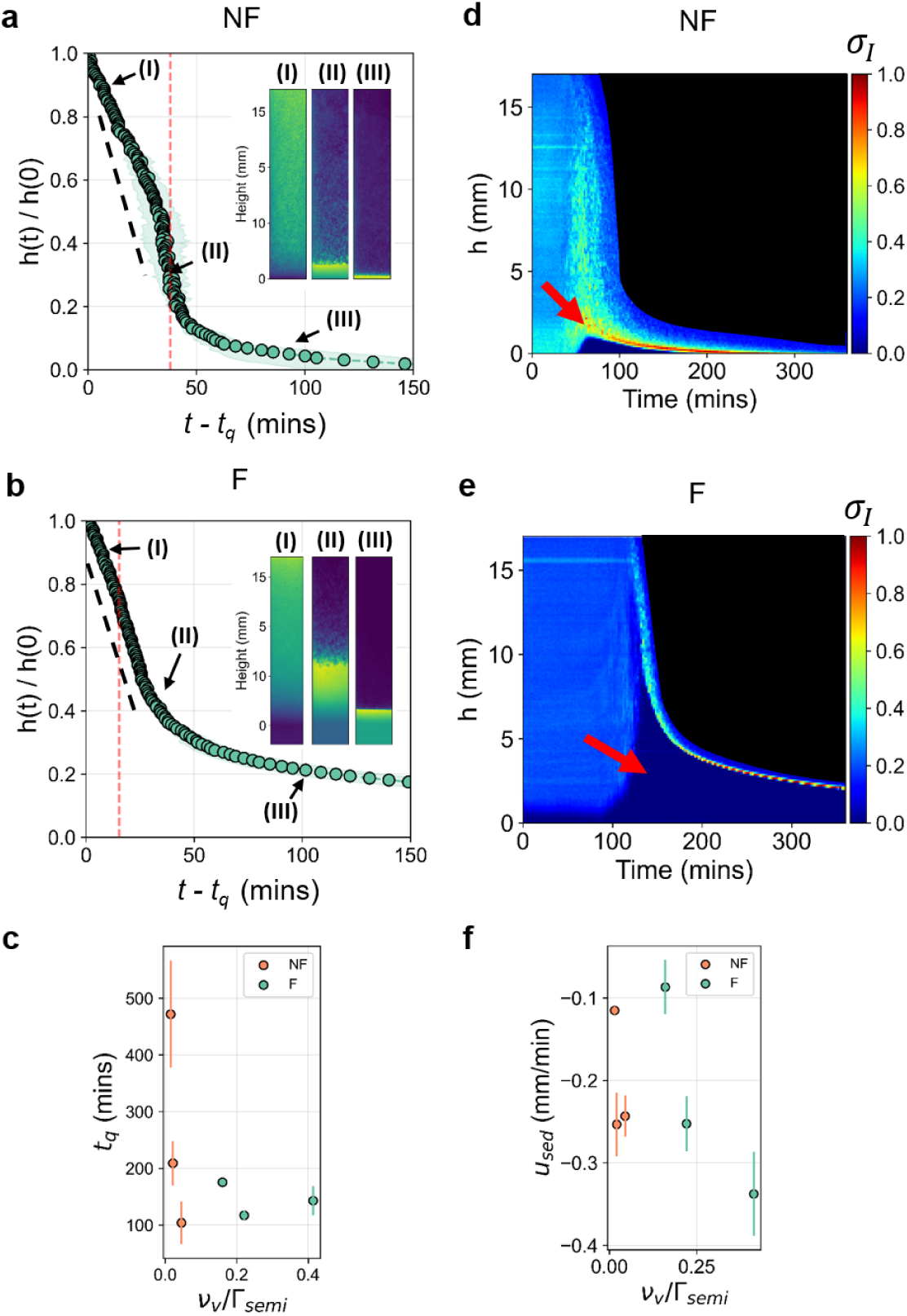
Bacterial suspensions display collapsing gel behaviour upon phase separation. (**a, b**) Normalised height, *h*(*t*)/*h*(0), as a function of time for the highest respective number density of NF and F bacterial suspensions after an initial quiescence time *t_q_* (stage 1). *h*(0) is the cuvette height. The curves display a linear relationship (I, black dashed line, stage 2) with time before the meniscus is formed (II, red dashed line). The meniscus collapsed following a stretched exponential relationship with time (III, stage 3). Insets (I, II, and III) are representative micrographs of the different stages. (**c**) The quiescence time, *t_q_*, is plotted as a function of respective number densities for both the NF and F bacterial suspensions. (**d, e**) Dependence of the normalised standard deviation in intensity *σ_I_*(*h, t*) with height, *h*, and time for the NF and F suspensions. The color bars indicate the scale of the normalised standard deviation in intensity, *σ_I_*. The gel point and collapse are highlighted with red arrows. The black region indicates the supernatant. (**f**) Sedimentation speed of the NF and F bacterial suspensions was measured from the linear portion of the collapse curve (stage 2, I to II) as a function of initial suspension number density. Displayed is the mean and standard deviation from at least 3 biological replicates.

While both phenotypes exhibit stages 1-3, the temporal dynamics of NF and F suspensions differ. First, the NF phenotype has a longer quiescent phase, *t_q_*, at the lowest number density, but this duration becomes comparable to that of the F cells at higher number densities (∼ 100 − 200 min, Fig. 2c). Second, during phase separation (stage 2), the suspension height decreases linearly with time at a velocity, *u*_sed_ (black dashed lines in Figs. 2a, b). The NF phenotype displays no clear dependency of *u_sed_* on the number density; however, the F phenotype displays a clear inverse relationship (Fig. 2f). Third, the transition between phase separation and compaction (stages 2 and 3, red dashed line in Figs. 2a, b) occurs at different heights for the NF and F systems, with the F phenotype reaching a significantly higher transition height (12.42 mm ±0.27 vs 3.42 mm ±0.30; fig. S2a). The onset of the compaction regime (stage 3) coincides with the emergence of a well-defined interface (insets (II) in Figs. 2a, b). The interface height follows a stretched exponential decay, indicating that the system has become gel-like (fig. S2b and (*39*)). Consequently, we define the beginning of the compaction regime (stage 3) as the “gel point”. From this point we see that NF suspensions develop networks that are less prone to compaction beyond the gel point, suggesting a higher mechanical rigidity in comparision to the F networks (fig. S2b).

Additional insight into the network assembly during sedimentation can be gained by quantifying the fluctuations in intensity, *σ_I_*, as a function of sample height, *h*, over time (Figs. 2 d, e, and Methods). This analysis shows that the nature of the interface between the dense and supernatant phases, as well as the fluctuations in intensity relating to bacterial density, are very different in the NF and F systems. In the NF phenotype, the interface is more diffuse (Fig. 2d), and there are stronger intensity fluctuations, suggesting that discrete clusters may be forming and sedimenting individually. In the F phenotype, instead, we observe a well-defined interface with lower intensity fluctuations, suggesting that a network has formed and is compacting during sedimentation (Fig. 2e). To understand the differences in the macroscale formation dynamics we examine the network formation on the microscale.

### Filamentation alters the microstructural and dynamical properties of cluster-cluster assembled bacterial networks

Next, we examine the microscale influence of filamentation on the dynamics of aggregation and subsequent microstructure of the cluster-cluster assembled bacterial networks. Here, we load NF and F cell suspensions into sealed glass wells (10 mm × 10 mm × 0.5 mm) and visualise them during sedimentation using optical microscopy (see Methods). We observe that *C. denitrificans* assembles into space-spanning networks on the surface of the capillary (Fig. 3a, b, top panels), which are branched and have a fractal-like structure (fig. S3).

**Figure 3:**
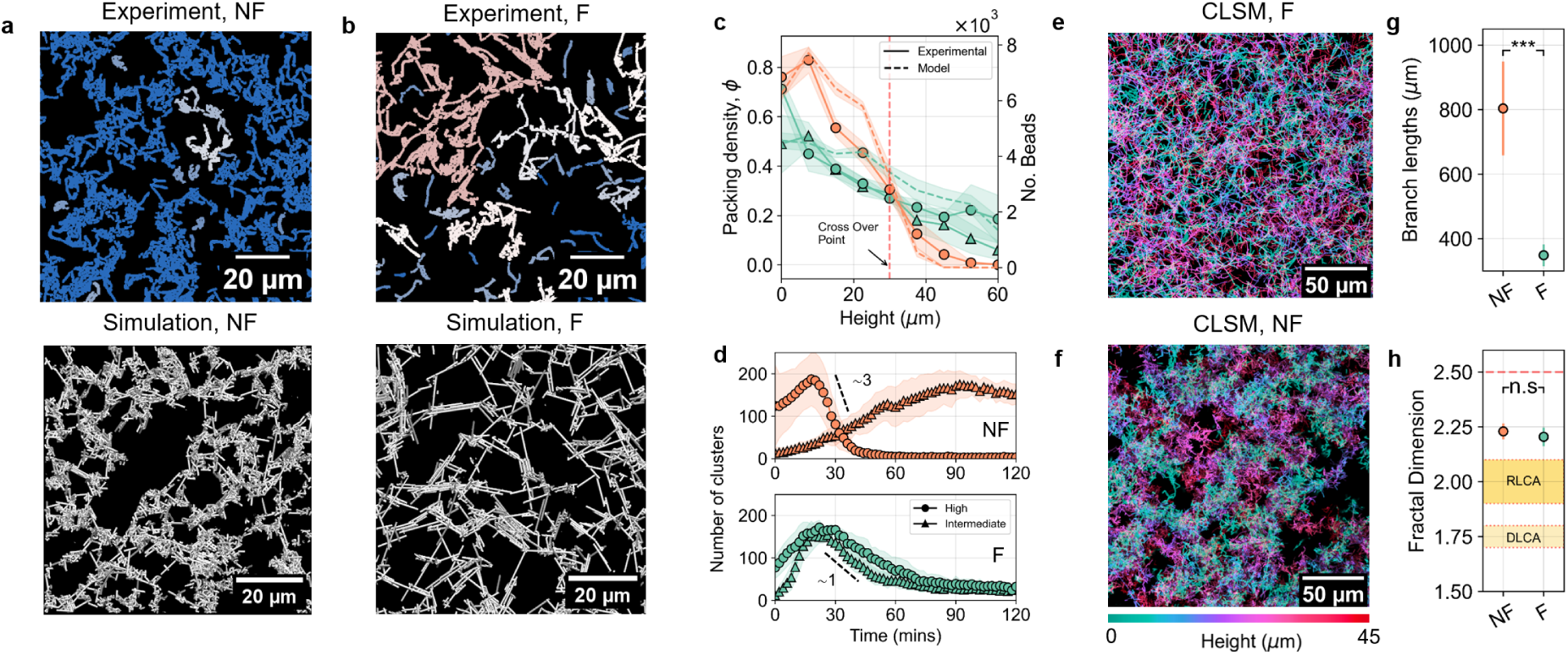
*C.denitrificans* forms bioflocculated gel-like structures on the microscale. (**a, b**) Representative micrographs of the NF and F bacterial networks acquired 30 *µ*m above the bottom surface in the experiments and (*>* 10*µ*m) in the simulations. The connected clusters in the experimentally obtained micrograph are indicated by the color coding. (**c**) Packing fraction dependence upon the height of the bacterial suspension. The NF phenotype is indicated in orange, and the F phenotype is indicated in green. Experimental data is indicated by the solid lines, while the simulation results are represented by the dashed lines of the respective phenotype colors. The red dashed line indicates the cross-over point at which clustering data were analyzed. (**d**) Respective cluster numbers for the NF and F phenotypes at intermediate and high initial cell number densities. (**e, f**) Z projections from CLSM volume scans of the F and NF networks, respectively, the false color representation of the slice height from the bottom of the networks. (**g**) Branch lengths of the bacterial gels were calculated from CLSM images taken of high-number density bacterial networks formed after 2 hours. (**h**) The fractal dimension of the bacterial networks formed after 2 hours; featured are the reaction-limited aggregation (RLA) and diffusion-limited aggregation (DLA) regions. Plotted are the mean and the standard deviation from at least 3 biological replicates, each with at least 3 different fields of view. Paired t-tests were performed for statistical analyses. n.s denotes non-significant, *** denotes P *<* 0.001.

The microstructure of these bacterial networks strongly depends on bacterial concentration and the filamentation phenotype (Fig. 3a-c). At the lowest number densities, both the NF and F cells settle on the bottom surface as quasi-2-D clusters (fig. S4). In this case, the settling rate is independent of filamentation (fig. S5a, b). At intermediate number densities, the F phenotype significantly accelerates the rate of network formation (fig. S5c, d). These settled structures form 3D space spanning networks, where the packing fraction decreases as a function of height, with the filamented networks plateaued at *ϕ* ≈ 0.4 after ≈ 60 minutes. This trend is more pronounced in the NF phenotype, as indicated by the steeper slope in Fig. 3c (red dashed line), which shows a higher concentration of cells at the base and a lower concentration at the top.

The assembly dynamics of the F and NF networks differ significantly. The assembly can be quantified by measuring the time evolution of the number of clusters, through time and height, from the brightfield experiments (Fig. 3d.). To allow for a valid comparison, we perform this analysis at the height of 30 *µ*m in the sedimentation profile, where the final packing fraction of the two systems is equal (cross-over point in Fig. 3c). There are two distinct phases of network formation; initially, cells collide and stick together to form small clusters; subsequently, clusters coarsen through collisions and grow into larger ones, eventually percolating, forming one (or very few) connected cluster(s) in a steady state. The transition between cluster formation and coarsening is marked by a peak in the number of clusters, which occurs after ∼30-minutes in the F phenotype at both intermediate and high cell concentration (Fig. 3d, lower panel). However, for the NF phenotype at intermediate concentration, coarsening occurs ∼ 60 min later, suggesting that the dynamics of the formation of the NF networks are more strongly dependent on cell concentration, consistent with the observed macroscale dependence on *t_q_*. Furthermore, filamentation results in a reduced exponent during the coarsening stage, with percolation occurring much more rapidly in the NF phenotype (exponent of −3) compared to the F phenotype (exponent of −1) at high concentration (Fig. 3d). Similar dynamics has been described in previous works and attributed to an almost restricted 2D aggregation (*40,41*). Specifically, the rate at which clusters percolate depends on the interaction between already settled and incoming clusters. The NF morphology has comparably larger voids between clusters, providing more free space for incoming clusters to settle into, thereby accelerating in-plane percolation. Qualitatively, the NF system coarsens and gels more quickly because it has greater connectivity, as indicated by the growth in the largest cluster size over time (fig. S6).

To probe the 3D microstructure of the networks, we use confocal laser scanning microscopy (CLSM) to image the bacterial network volume (Fig. 3e, f and fig. S7). We compute the total branch lengths of the composing clusters and their fractal dimension (Fig. 3g, h). Bacterial clusters are, on average, significantly shorter in the F phenotype (Fig. 3e), while the network fractal dimension is consistent across the phenotypes (F networks; 2.20 ±0.04 vs NF networks 2.23 ±0.03 (Fig. 3g). The fractal dimensions of the bacterial networks align with those of other anisotropic gel systems, such as clay platelets and colloidal discoid gels (*42, 43*). Anisotropic systems have frequently been reported with higher fractal dimensions than isotropic gels, attributed to anisotropic excluded volume effects (*44*).

We then monitored the post-assembly dynamics of the surface-bound bacterial networks as they developed into biofilms. To do this we form bacterial networks within microfluidic channels, where a creeping flow ensures a constant nutrient supply, and image the network development for 10 hours. The foundational network structures increase in the area fraction over time, reflecting system densification as the network grows and matures into biofilm. Furthermore, fluctuations in the motion of the network progressively quench, indicating the complete structural arrest expected in mature biofilms (fig. S9).

To further understand the dynamics observed in our experiments, in particular, the emergence of the network architecture, we perform Langevin dynamics simulations (see Methods). These simulations are initialised with a dilute polydisperse suspension of rigid rods with length distribution matching that of the experimental bacterial system. Each rod is modeled as a straight chain of connected beads, and cohesion between cells is simulated by allowing beads on one rod to interact with beads on other rods during sedimentation. This approach is sufficient for capturing the self-assembly of a gel-like structure for a rod density that is large enough. We observe that the resulting gels depend strongly on the average rod length (i.e. the experimental filamentation phenotype), influencing both the overall morphology and the critical density for gel formation (or percolation threshold). Remarkably, our simulations also yield a density profile, which is in near-quantitative agreement with microscopy experiments (Fig. 3c). The weaker but linear relationship (shallower slope Fig. 3c) the F phenotype exhibits points to the presence of a space-spanning network, which, contrary to the NF phenotype, also forms at intermediate cell concentration (Fig. 3c, triangles).

### Filamentation shapes the mechanical response and yielding modes of bacterial networks

In colloidal gels, the cohesion between the constituent particles confers structural stability and the emergence of elasticity, both traits we hypothsise the bacterial networks to display. To measure the microscale mechanical response of the bacterial networks, we invert the capillaries, containing bacterial networks on the capillary floor, subjecting the networks to gravitational stress. The network deformation is then tracked for 2 hours and quantified using the normalised packing density, *ϕ*(*t*)*/ϕ*_0_, where *ϕ*(*t*) is the packing density at time, *t*, and *ϕ*_0_ is the packing density at the beginning of the reversal experiment. The networks either yield or remain intact under gravitational stress (Fig. 4a, b), with yielding occurring in three distinct ways depending on filamentation phenotype. First, the networks can undergo viscous-like cluster-cluster deconstruction, where individual clusters gradually break away from the collective structure (type I yielding; movie. S1). Second, we observe delamination of the entire network structure, originating at the base layer (type II yielding). Third, in some cases, we also see elastic-like fragmented yielding, where subregions of the network fracture and break away from neighbouring regions which remain intact and surface attached, indicating viscoelastic solid behaviour (type III yielding; movie. S2). Networks that do not yield instead relax slowly with time until a plateau is reached (movie. S3).

**Figure 4:**
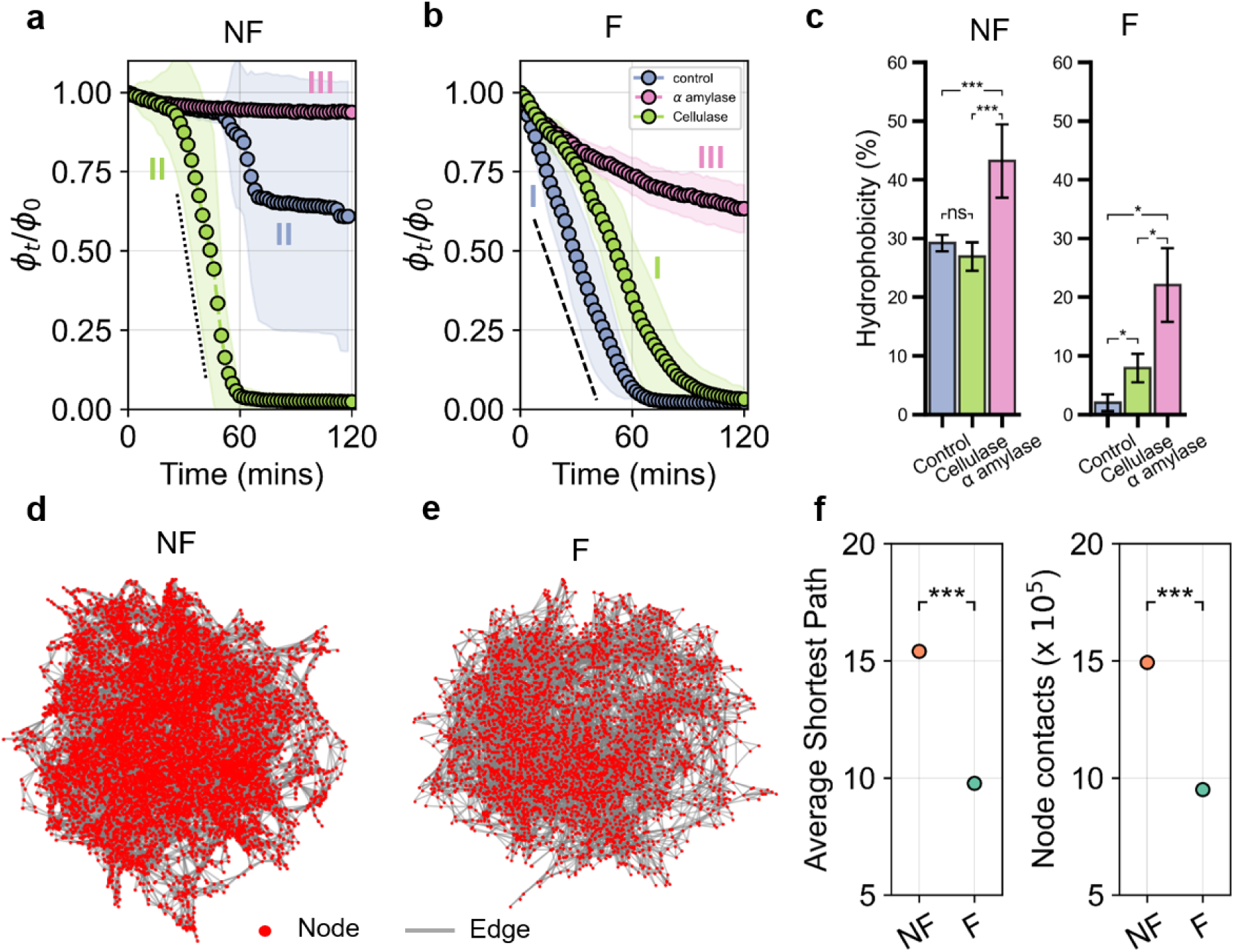
Micro-mechanical peeling, connectivity, and network strength vary as a function of filamentation and cell surface hydrophobicity. (**a, b**) Normalised change in packing density, *ϕ_t_/ϕ*_0_, from 30 *µ*m above the base of the bacterial networks for F and NF cells, respectively. The data presented are the mean and standard deviation from at least 3 biological replicates, each with 5 different fields of view. (**c**) Cell surface hydrophobicity is influenced by enzymatic treatments. The cell surface hydrophobicity of F and NF cell suspensions treated with *α*-amylase and cellulase enzymes. The controls are untreated cells. (**d, e**) The average network path and the number of node contacts are calculated from the network graphs. The data presented are the average and standard deviation from 3 simulation runs. Paired t-tests were performed for statistical analyses. n.s denotes non-significant, * denotes P *<* 0.05, *** denotes P *<* 0.001, **** denotes P *<* 0.0001. (**f**) Network analysis of F (d) and NF (e) cell length distributions from the model. The red dots indicate nodes, and the grey lines indicate the edges of the network graph.

Networks assembled from F cells yield through type I cluster-wise detachment with a rate of ∼ −0.03 min^−1^ over a 60-minute period (Fig. 4b, dashed line). Upon interrogation of the final structure on the capillary ceiling, we see a layer of surface-attached cells and tufts (fig. S9). In contrast, the NF networks initially remain intact; however, after ≈ 60 minutes, a subset of the networks (33%, fig. S10) experienced an elastic-like fragmented type III yielding from the ceiling (Fig. 4a, dashed line). These differences suggest that the F phenotype induces the formation of bacterial networks with mechanics more viscous-like than that of the NF phenotype.

The differences in mechanical behaviour due to the induction of filamentation can be explained by network topology. The elasticity of colloidal gels is known to be mediated by the number of contacts between discrete clusters and the strength of interparticle interaction (*21*). To gain insight into the number of contacts in the bacterial networks, we perform a connectivity analysis on the final network configurations generated in our Langevin dynamics simulations (see Methods). The resulting contact maps reveal significant topological differences with the F networks having fewer contacts (9.50 × 10^5^ ± 0.06 vs 14.93 × 10^5^ ± 0.06) (Fig. 4d, e) and shorter network path lengths (9.77±0.28 vs 15.40±0.18) (Fig. 4f), in agreement with the CLSM branch length analysis (Fig.3g). Additionally, we quantify the cell surface hydrophobicity to assess the interaction strength between the cells, offering an insight into the cell surface electrostatic effects (see Methods). NF cells are significantly more hydrophobic than the F cells (29.17 %±2.97 vs 1.99 %±1.41) (Fig. 4c). Together, these data suggest that filamentous bacteria are more hydrophilic and assemble into networks with fewer contacts that result in a bias towards more viscous-like yielding dynamics.

### Enzymatic treatment of bacterial gels tunes network strength

Having established the baseline mechanical response of bacterial networks, we sought to deduce the influence of cell surface biopolymers on the underlying mechanics. We pre-treated cells with the enzymes *α*-amylase and cellulase to specifically hydrolyse *α*-1,4 and *β*-1,4 polysaccharide linkages, respectively (*45*). Phase contrast microscopy reveals that treatment with *α*-amylase degrades the polysaccharide components on the cell envelope of *C. denitrificans*, causing a significant loss in cell viability (figs. S1 and S11). Treatment with cellulase, on the other hand, has little effect on the integrity of the cell envelope and cell viability. Regardless of phenotype, treatment with *α*-amylase significantly increases cell hydrophobicity, particularly against the F cells (from 1.99%±1.41 to 22.01 %±6.26 for the F cells compared to 29.17%±2.97 to 43.15 %±0.42 for NF cells). Cellulase treatment significantly increased the hydrophobicity of F cells (7.90 %±2.42; Fig. 4c).

Enzymatic modification of the cell surface alters the mechanical response of bacterial networks to gravitational stress. First, the F and NF cells pre-treated with *α*-amylase produce networks that transition from type I and type III yielding behaviours, respectively, to no yielding (*ϕ_t_/ϕ*_0_ = 0.95 ± 0.07), and a purely extensional deformation mode (*ϕ_t_/ϕ*_0_ = 0.63±0.07), indicating an increase in network strength, that also coincides with a reduction in cell viability (Fig. 4a, b). Second, in the cellulase-treated F cells, we observe a delay in the onset of type I yielding (≈ 25-minutes) in comparison to the control. In contrast, in cellulase-treated NF networks, the lag time preceding type II yielding (delamination) reduces from 60-minutes to 30-minutes (Fig. 4a). Together, these yielding behaviours suggests that surface/network adhesion, rather than the intra-network cohesion, induces mechanical failure, as the networks remain otherwise intact during delamination. Furthermore, the dynamics of the NF networks are more symptomatic of elastic, solid-like yielding.

### Filamentation promotes an increasingly viscous mechanical response

To quantify the viscoelastic response of the NF and F bacterial networks, we conduct macroscale oscillatory shear rheology using a rotational rheometer. We measure the F and NF networks using the volume fraction range found in the microscopy experiments (*ϕ* = 0.1 - 0.4). Both networks displayed a solid-like, elastically dominated response within the linear viscoelastic regime, where *G*^′^ *> G*^′′^, regardless of the cell phenotype, thus confirming that the bacterial networks behave as gels. When plotted against their respective volume fraction, *ϕ*, the F networks have a higher linear elastic, G’ and viscous moduli, G” (Fig. 5a, b). For example, at a volume fraction of *ϕ* = 0.4, the F network was significantly more elastic than the NF (9.48 ±1.17 Pa vs 5.17±0.44 Pa). This increased elasticity is attributed to the greater network homogeneity in the filamented networks, as observed in the CLSM imaging (Fig. 3e), similar to findings in anisotropic colloidal gels (*46*).

**Figure 5:**
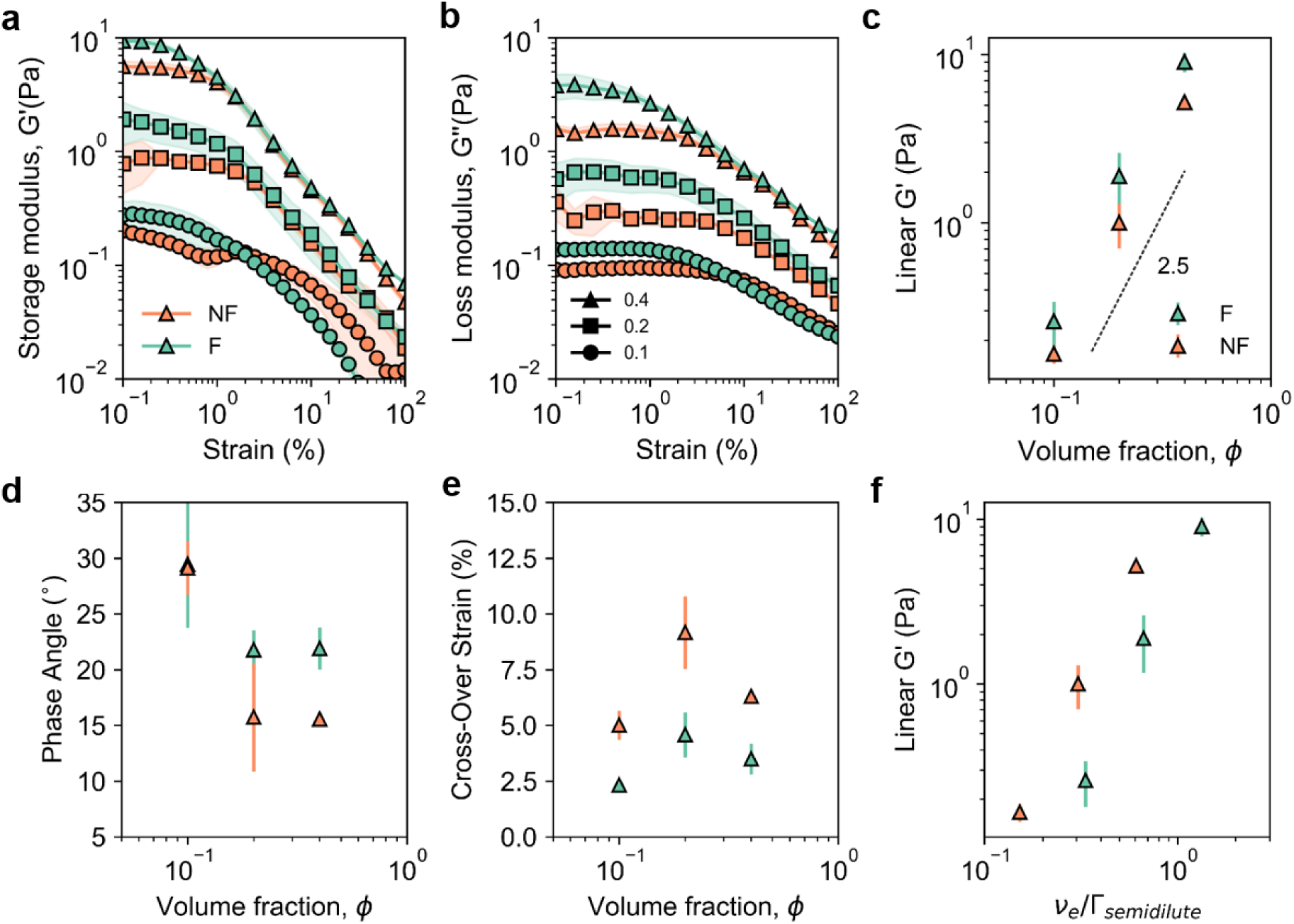
Bulk rheology of the bacterial networks. (**a, b**) Storage (G’) and loss (G”) modulus for the F and NF networks at volume fractions *ϕ* of 0.1, 0.2, and 0.4 from the linear to nonlinear range. (**c**) Scaling of the linear storage modulus (G’) taken from the linear range of the amplitude sweeps against the respective packing fractions. (**d**) The phase angle of the networks is calculated from the linear viscoelastic range. (**e**) The cross-over strain of the N and NF networks, defined as the strain value where G’ ¡ G”.(**f**) The linear storage modulus (G’) as a function of the NF and F number density normalised by the semi-dilute limit for the respective ^systems, Γ^*semi*−*dilute*

The plateau elastic moduli of both networks displays a power law dependence on volume fraction, scaling as *∝ ϕ*^2.5^. (Fig. 5c). This scaling is similar to that observed in attractive rod (*∝ ϕ*^2.3^) and discoid gels (*∝ ϕ*^2.5^) (*42, 47*). Both network phenotypes exhibit a yield strain of ≈ 1%. Upon yielding, the elastic and viscous moduli of the two networks converge. Interestingly, however, the phase angle of the NF bacterial networks is consistently lower than their filamented counterpart, indicating increased solid-like dynamics in the absence of filamentation (Fig. 5d). This result is consistent with the microscale yielding observations, where the NF networks display a more-pronounced solid-like yielding mode (Type II, fracture). Additionally, the crossover strain, indicative of the fluidisation transition, increases in the NF case at the two highest volume fractions tested (Fig. 5e), and the cross-over stress is higher in the NF case (*σ_c_* = 0.068±0.004 Pa vs *σ_c_* = 0.058±0.017 Pa). Finally, considering the polydisperse cell lengths in the system, we compare the respective number densities of the networks against their theoretical semi-dilute limits (Fig. 5f). After this correction, normalizing cell overlap reveal that the NF networks have a higher comparable elasticity.

## Discussion and Conclusion

Here, we quantified the dynamics of bacterial aggregation in a system that assembles into settled networks via cluster-cluster aggregation. By controlling the induction of cellular filamentation, we explored the influence of cell length and cell surface hydrophobicity on the formation of the space-spanning networks and their structure-function properties, revealing their gel-like dynamics. Using Langevin dynamics simulations that account for the polydispersity in cell lengths, we elucidated the microstructural properties of the assembled bacterial networks using simple interaction potentials. Notably, the cluster-cluster assembled bacterial networks bared a volume fraction- storage modulus dependence analogous to attractive colloidal rod systems.

In biofilms, filamentation is known to bridge microscale colonies in patchy environments, while in fluctuating flow, it enhanced early colonisation (*13, 48, 49*). Our findings suggest that in the context of aggregate formation, filamentation enhances the effective number density reducing the cell concentration required to induce phase separation and the formation of clustercluster aggregated bacterial networks. In non-interacting colloidal hard rod systems, the fat-tails of the log-normal rod length distribution move the isotropic-nematic phase boundary to lower density (*50*). Therefore, it is plausible that the percolation boundary is reduced by filamentation. Although filamentation induced network formation at lower respective cell numbers, they are mechanically weaker. Our microstructural analysis indicates F settled networks are more homogeneous but have fewer connections than NF cell networks. Upon surface settling, the assembled network then densifies with time and cell growth into a biofilm structure (fig. S9).

The macrorheological scaling behaviour of cluster-assembled bacterial networks follows a similar exponent to that of anisotropic colloidal gels of comparable aspect ratios (*42, 46*). Our results indicate that filamentation reduces the solid-like response of the networks, inducing microscale cluster-cluster breakdown under gravitational stress and an increased phase angle when tested at the macroscale. Given that purely physical parameters govern these phenomena, we suggest that the basic physical principles derived from the *C. denitrificans* system may be transferable to other bacterial species. However, species-specific variations in cell length distribution and interaction strength, dependent on cell surface biopolymers, are likely to quantitatively affect the assembled structure and sedimentation dynamics. Our findings also suggest that cell surface hydrophobicity plays a significant role in modifying the yielding dynamics and network cohesion.

Our work bears relevance to real-world environments such as bioreactors and wastewater treatment plants, where bacterial flocculation and settling are integral parts of the process. In addition to the ecological significance of understanding cluster-cluster biofilm assembly, this research offers an intriguing avenue toward fabricating living material network structures guided by sedimentation. In particular, a better understanding of the role of cellular morphology could help optimize the function and material properties of bacterial gels for their incorporation within bioreactor systems.

## Methods and materials

### Bacterial Culture

Experiments were performed using *Comammonas denitrificans* 123 (ATCC 700936). *Comammonas denitrificans* suspensions were prepared by inoculating 5 ml tryptic soy broth (TSB) with cells from glycerol cryo stocks and incubated at 30^◦^C with 180 rpm shaking. Dissolved oxygen conditions were controlled by modulating the head space in the culture vessels. To create O_2_ depleted conditions 14 ml culture tubes were used, which provided a 1.8× headspace with respect to the liquid volume. O_2_+ conditions were created by using 50 ml falcon tubes, which provided a 9× headspace with respect to the liquid volume. To screen the incubation conditions to induce cell elongation, overnight cultures were diluted 1000:1 in the O_2_-limited and O_2_-replete culture vessels. Cells were then incubated with and without shaking (180 rpm) at 21^◦^C, 30^◦^C, and 37^◦^C for 24 hours. Three biological replicates were performed for each condition.

### Cell length distributions

To measure cell lengths, cell cultures were grown to OD = 1.0. 2 *µ*L of culture was transferred to a 1.5 % agarose pad, which was covered with a No 1.5 coverslip. Imaging was performed using a Nikon Ti inverted microscope (Nikon, Japan) with a digital camera (Zyla sCMOS, Andor, UK). Bacterial cells were imaged in phase contrast mode using a 63× oil immersion objective. Three biological replicates were performed, and up to 15 fields of view (147 *µ*m x 147 *µ*m) were acquired per biological replicate. The cell length was calculated from the images by applying Otsu binarisation and then quantified using the skeletonization function in Fiji ImageJ. The images were padded, and cells at each frame’s edge were removed from the analysis.

### O_2_ concentration measurements

O_2_ concentration during cell culture was measured using an optical oxygen measurement system (PreSens Oxygen Meter, PreSens, Germany). Oxygen-sensitive probe spots adhered to the inside of the relevant culture vessels with manufacturer-provided adhesive. Vessels were then sterilised using isopropanol. The sensor was calibrated using oxygen-saturated water and zeroed with a solution of 300 mM sodium sulfite. Culture vessels were filled with 5 ml TSB media and inoculated at a 100:1 ratio from overnight cultures. Oxygen concentrations were measured every hour for 8 hours. For each condition, four biological replicates were used.

### Horizontal imaging setup

We determined the sedimentation behaviour and collapse of the meniscus using digital imaging. Samples were prepared to the relevant number density and homogenised using gentle vortexing before transfer into 2.5 mm × 10 mm × 20 mm plastic cuvettes for time-lapse observation at 21^◦^ C. The imaging platform comprised a CMOS camera (SVS-Vistek, Vision Systems LLC) with a 35 mm lens (HF3514V-2, Myutron). An LED panel provided illumination with a diffuser, and images were acquired in transmission mode. Images were acquired every 2 minutes for at least 300 minutes. To obtain the sedimentation behaviour and collapse of the meniscus, image analysis was performed. The sedimentation dynamics of each bacterial suspension were calculated by tracking the intensity profile through the height of the cuvette. The cuvette was split into horizontal regions of interest across the central 2 mm of the cuvette with a height of 5 pixels. The intensity in these regions was tracked in time; when the intensity dropped below a threshold value, the region was deemed clear of cells. The time taken for each region to be void of cells was measured for each region of interest, which enabled us to track the height of the suspension before a clear meniscus had formed. The quiescence time, *t_q_* was calculated as the time taken for the top region of interest of the cuvette to become void of cells. The sedimentation velocity, *u_sed_* was calculated as the gradient from the linear portion of the *h*(*t*) curve until the meniscus formed for each concentration and phenotype.

The emergence of the meniscus was characterised by a sharply contrasting interface between the supernatant and the sample. The emergence of the meniscus was calculated by tracking the vertical line profile of the central 2 mm region of each frame. A meniscus was deemed to have formed when the intensity gradient exceeded a threshold value corresponding to the interface between the supernatant and dense bacterial phase. The height of the meniscus interface was then tracked with time. To calculate the standard deviation in intensity with time, the raw images were split into regions of interest of 4 pixels high and across the central 2 mm of the cuvette. The standard deviation in intensity, *σ_I_* was then calculated for each of these regions across the observation period.

### Sedimentation and reversal experiments

For each experiment cell suspensions were grown to OD = 1.0, 1 ml of this culture was aliquoted into a 1.5 ml centrifuge tube. Cells were gently centrifuged for 90 seconds at 2700 rcf, the supernatant was discarded and cells were resuspended in 1 ml, 0.5 ml, and 0.25 ml of media. For experiments involving enzyme treatment, the cells were resuspended either 100 mg/ml *α*-amylase (Sigma Alderich, Switzerland) or 125 mg/ml cellulase (Sigma Alderich, Switzerland), re-suspended and incubated without shaking at 30 ^◦^C for 2 hours. Cells were then vortexed for 90 seconds to re-suspend and create a well-dispersed suspension. We ran control experiments to confirm that centrifugation and vortexing had a negligible effect on network formation.

Gel network formation through sedimentation was performed on an inverted microscope (magnification 20x + 1.5x zoom) with a piezostage (Nikon Ti, Nikon, Japan). The imaging cell (10 mm x 10 mm x 500 *µ*m) was created by layering 2 geneframes (Sigma Alderich, Switzerland). 54 *µ*L of polydisperse cell culture was pipetted into the imaging area and sealed using a No. 1.5 coverslip. The slide was immediately transferred to the microscope, and imaging commenced no later than 5 minutes after sample preparation. Imaging was performed using brightfield mode with Koeller illumination condition with the condenser 2/3 closed. Z stacks of 10 *µ*m increment from the bottom surface of the imaging area to 90 *µ*m into the imaging volume were acquired every 2 minutes for 120 minutes. 5 fields of view (441 *µ*m x 441 *µ*m) were acquired for each experimental condition, and at least 3 biological replicates were tested. Reversal experiments followed the same sedimentation procedures; however, after 2 hours of sedimentation, the imaging cell was inverted. Z stacks were then acquired from the top of the imaging arena to 90 *µ*m depth. Confocal scanning laser microscopy (Zeiss LSM 710, Zeiss, Germany) imaging was performed at 20x using the same procedure, with the exception being a staining step with Syto 9 (5*µ*M)(Sigma Alderich, Switzerland) before insertion into the capillary vessel. The bacterial networks were imaged at the end time point of each experiment through the sample height.

Raw images were processed using Fiji and a custom image analysis script using Python. Briefly, raw images were smoothed using a Sigma filter and binarised using a sub-pixel curvilinear structure extraction algorithm (*51*) (fig. S14). Binarised images were then eroded before a dilation step was applied to remove singular pixels. The packing density *ϕ* for each height and time point was calculated by 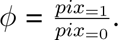 . The number of clusters was calculated using the *regionprops* function. For the CLSM stacks, the branch lengths and fractal dimensions were calculated using the *kimimao* package and the box-counting method, respectively (*52*).

### Rheometry

The viscoelastic properties of the bacterial networks were measured using a rotational rheometer (Anton Paar). Bacterial cultures were harvested after 10 hours for the NF case and 24hours for the F phenotype using the aforementioned respective culture methods. Cell cultures were centrifuged and the supernatant was discarded. Cells were pooled from 5ml of culture volume to yield 3 - 5 biological replicates per volume fraction and filamentation phenotype. Cells were then spun down, supernatent was discarded and the mass of cells was measured using a high precision balance. Based on the mass, culture media was added at the appropriate volume to obtain volume fractions of 0.1, 0.2, and 0.4. The cell suspensions (350 *µ*m) were pipetted onto a roughened bottom plate geometry. A 20 mm roughened cone and plate geometry with a gap height of 150*µ*m was used. The sample was left to equilibrate for 5 minutes before an amplitude sweep was performed (*γ* = 0.01 - 100%) at a frequency of 0.1592 rads^−1^.

### Cell surface hydrophobicity measurement

We measured the comparative cell surface hydrophobicity (CSH) of the cell suspensions by performing the microbial adhesion to hydrocarbon (MATH) assay (*53*). Bacterial cultures were grown to an OD of 1 as previously described. Upon sampling, 1 ml of the culture was centrifuged (2700 rcf) for 2 minutes and cleaned 3 times with fresh media. Resuspended cells were then diluted to an *OD*_600_ of 0.5 in a 5 ml test tube. 4 ml of cell suspension was mixed with 1 ml of hexadecane (Sigma Aldrich) and vortexed at full speed for 2 minutes. Mixed suspensions were then left to phase separate for 30 minutes.

After a syringe with a needle tip was used to collect 1 ml of cell suspension from the aqueous phase. The OD of this suspension was then measured. The percentage hydrophobicity was calculated as 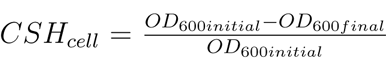. Low % values indicate hydrophilicity, whilst high % values indicate hydrophobicity.

### Effective Number density calculations

The effective number density, *ν_e_*, of the cellular suspensions was calculated using the measured cell length distributions for the NF and F cells and their respective cell concentrations (cells ml^−1^) obtained from the flow cytometry measurements (CytoFLEX, Beckman Coulter). The number density for rods with a defined length can be calculated as 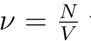 where *N* is the number of rods and *V* is the sample volume. However, for polydisperse suspensions of rods with a length distribution, the effective number density is calculated as 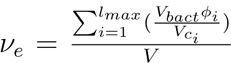 (54). In the equation, *l_max_* is the longest cell length, *V_bact_* is the volume of cells in the sample, *ϕ_i_* is the volume fraction of cells with length *i*, and *V_ci_* is the volume of a single cell with length *i*. We estimate *V_ci_* using a spherocylindrical geometry. The dilute boundary was calculated as 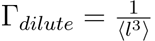.

### Langevin simulations

We model our non-growing bacterial cells as rigid straight chains of connected beads, with neighbouring beads separated by a distance *σ*. The length of each rod, in units of beads (or *σ*), is drawn from a log-normal distribution corresponding to the “filamentous” (mean = 1.848*σ*, sd = 0.574*σ*) and “nonfilamentous” (mean = 1.050*σ*, sd = 0.469*σ*) phenotypes in Fig. 1e.

Beads interact with beads on other rods via a cut-and-shifted Lennard-Jones (LJ) potential,

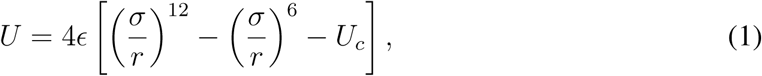

where *ɛ* governs the strength of the interaction, *σ* is the bead diameter, and *U_c_* = (*σ/r_c_*)^12^ −(*σ/r_c_*)^6^ ensures that the potential is equal to zero at the cut-off distance *r_c_*. For equilibrium runs, *r_c_* = 2^1^*^/^*^6^*σ*, corresponding to the Weeks-Chandler-Anderson (WCA) potential. Otherwise, we set *ɛ* = 50 and *r_c_* = 1.4*σ* in order to model strongly attractive and short-ranged interactions between the beads. Here, we set our energy in units of *k_B_T*, where *k_B_* is the Boltzmann constant and *T* is the temperature. Furthermore, given that we are interested in bacteria, we set our characteristic lengthscale *σ* to be 1*µ*m.

We use the Large-scale Atomic/Molecular Massively Parallel Simulator (LAMMPS) (*55*) to perform Langevin Dynamics (LD) simulations of our sedimenting colloidal rods. The time evolution of each bead is governed by the Langevin equation,

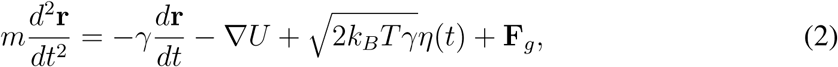

where *U* is the potential energy of the bead resulting from the sum of interactions with all other beads in the system, *m* is the mass of the bead, *γ* is the friction coefficient, **F***_g_* is the force due to gravity, and *η* is a white noise term with zero mean and variance given by

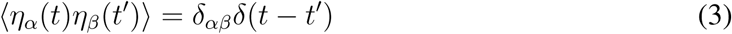

along each Cartesian component. This equation is integrated using a velocity-Verlet scheme.

In our simulations, we consider the time *τ_D_* for a bead to diffuse its own length, *σ*, as the natural unit of time, which is related to the diffusion coefficient *D* of a colloidal particle through 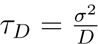. Using the Stokes-Einstein relation 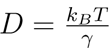, we can relate the diffusion coefficient *D* to the friction coefficient *γ*, which in turn relates to the dynamic viscosity *µ* through *γ* = 3*πµσ*.

We simulated our bacterial suspensions at a volume fraction of 0.015. Initial configurations for any given system were set up by filling a simulation box (80 × 80 × 160*σ*^3^) with rods drawn from either the F or NF distribution. Rods were constructed by placing the first bead of each rod at a random location in the box and sequentially adding extra beads along the *x* direction at distances *σ* until the length of the rod matched that drawn from the distribution.

A 3-step equilibration of this initial configuration was then performed. In the first step, particle overlap resulting from the initial configuration was removed by “relaxing” the system using a soft cosine potential for 5 × 10^2^*τ* . Then, in the second step, the system was allowed to “relax” further by switching to a WCA potential for 2×10^2^*τ* . Finally, we increased the timestep from 5 × 10^−4^ to 5 × 10^−3^*τ* for a further 10^4^*τ* to distribute the bacteria evenly throughout the simulation box.

For both F and NF suspensions, four different initial configurations (resulting from different input random seeds) were relaxed in this three-step process to generate four equilibrated configurations. These four equilibrated configurations were then used as the starting configuration for the sedimentation simulations of duration 10^4^*τ* . To simulate gravity, a downward force (**F***_g_*) of -0.1 *k_B_T/σ* was applied to each bead. Thus four simulations were performed, each with a different starting configuration and random seed for the Langevin noise.

To quantify the network features from simulation data, we proceed as follows. First, a network is constructed from a configuration (simulation snapshot) by identifying rods as nodes, and by adding an edge between each pair of rods which is below a threshold distance (equal to 1.2*σ*, note the distance between two rods is computed as the minimum between the pairwise distance of any pair of beads belonging to the two different rods). Then, we characterise the connectivity of each network by computing: (i) the total number of connections (or edges), and (ii) the average shortest path between a node pair. The analysis has been performed by using the Python library NetworkX (*56*). Values reported in the main text are then averaged over different configurations for each condition.

## Acknowledgments

The authors would like to thank Dr. Jonasz Slomka and Dr. Chris Brackley for valuable discussions. The authors acknowledge financial support from SNSF PRIMA grant 179834 (to E.S.), MSCA individual fellowship grant 101033169 (to S.C.) and NBIC/BBSRC/UKRI grant No. BB/R012415/1 (to G.M).

## Supplementary materials

Materials and Methods

Figs. S1 to S14

References

## Other Supplementary Materials for this manuscript include the following

Movies. S1 to S3

