## Supplementary information for "The role of cellular filamentation in bacterial aggregation and cluster-cluster assembly"

### Supplementary Materials for:
### The role of cellular filamentation in bacterial aggregation and cluster-cluster assembly

Samuel Charlton[1], Gavin Melaugh[2,3], Davide Marenduzzo[2], Cait MacPhee[3], and Eleonora Secchi[1,*]

[1]Institute of Environmental Engineering, Department of Civil, Environmental and Geomatic Engineering, ETH Zürich, Zürich, 8093, Switzerland
[2]SUPA, School of Physics and Astronomy, University of Edinburgh, Edinburgh, EH9 3FD, UK
[3]School of Engineering, University of Edinburgh, Edinburgh, EH9 3JL, UK

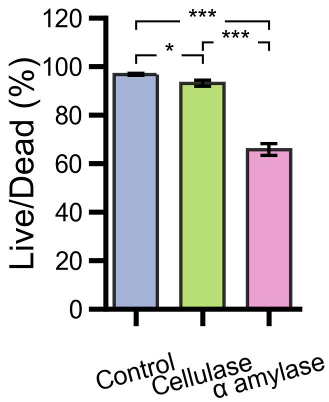


**Supplementary figure 1: Live dead assay on *Comamonas denitrificans* after enzyme exposure.** Live/ dead % for cell suspensions after 2 hr enzyme exposure. Shown are the mean and standard deviation from 3 biological replicates. * denotes P < 0.05, *** denotes P < 0.001.

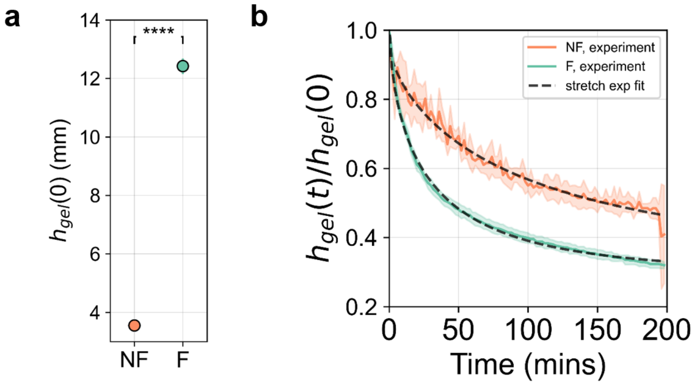


**Supplementary figure 2: Gel point height for the filamented (F) and non-filamented (NF) cells and the collapse dynamic.** (**a**) The gel point $h_{gel}(0)$ was determined when a clear interface was formed between the supernatant and concentrated cell phase during the macroscale sedimentation tests for the highest number densities for each phenotype. Shown are the mean and standard deviation from at least 3 biological replicates. (**b**) The gel interface $h_{gel}(t)$, was tracked with time for the NF and F cell suspensions and normalised by the initial height at which the gel was deemed formed, $h_{gel}(0)$. The collapsing dynamic followed a stretched exponential fit (black dashed line). Significance was tested using the student t-test **** denotes P < 0.0001.

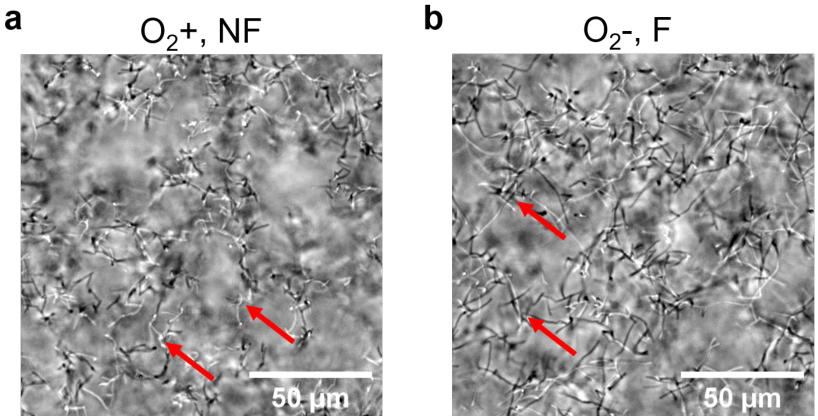


**Supplementary figure 3: Representative phase brightfield images of the bacterial networks.** Shown are the $O_2$+ (**a**), NF and the $O_2$- (**b**), F cell networks formed at the in highest respective number densities. The red arrows indicate cell/cell contracts of both tip-tip and tip-body. The images shown are taken 30 $\mu$m above the bottom surface of the capillary.

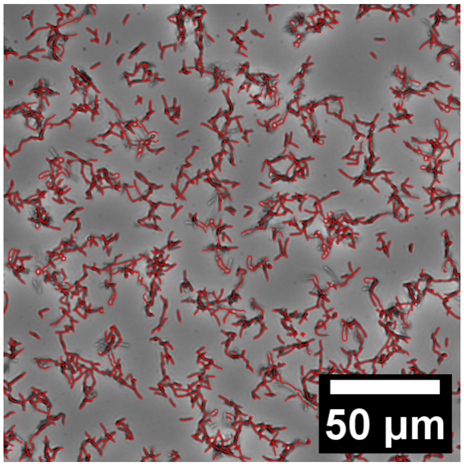


**Supplementary figure 4: Representative micrograph of quasi-2D cell clusters.** Representative micrograph of the flocculated network structure formed by the low number density NF phenotype upon sedimentation after two hours; the slice presented is taken at the glass capillaries bottom surface. Red lines indicate connected network structures.

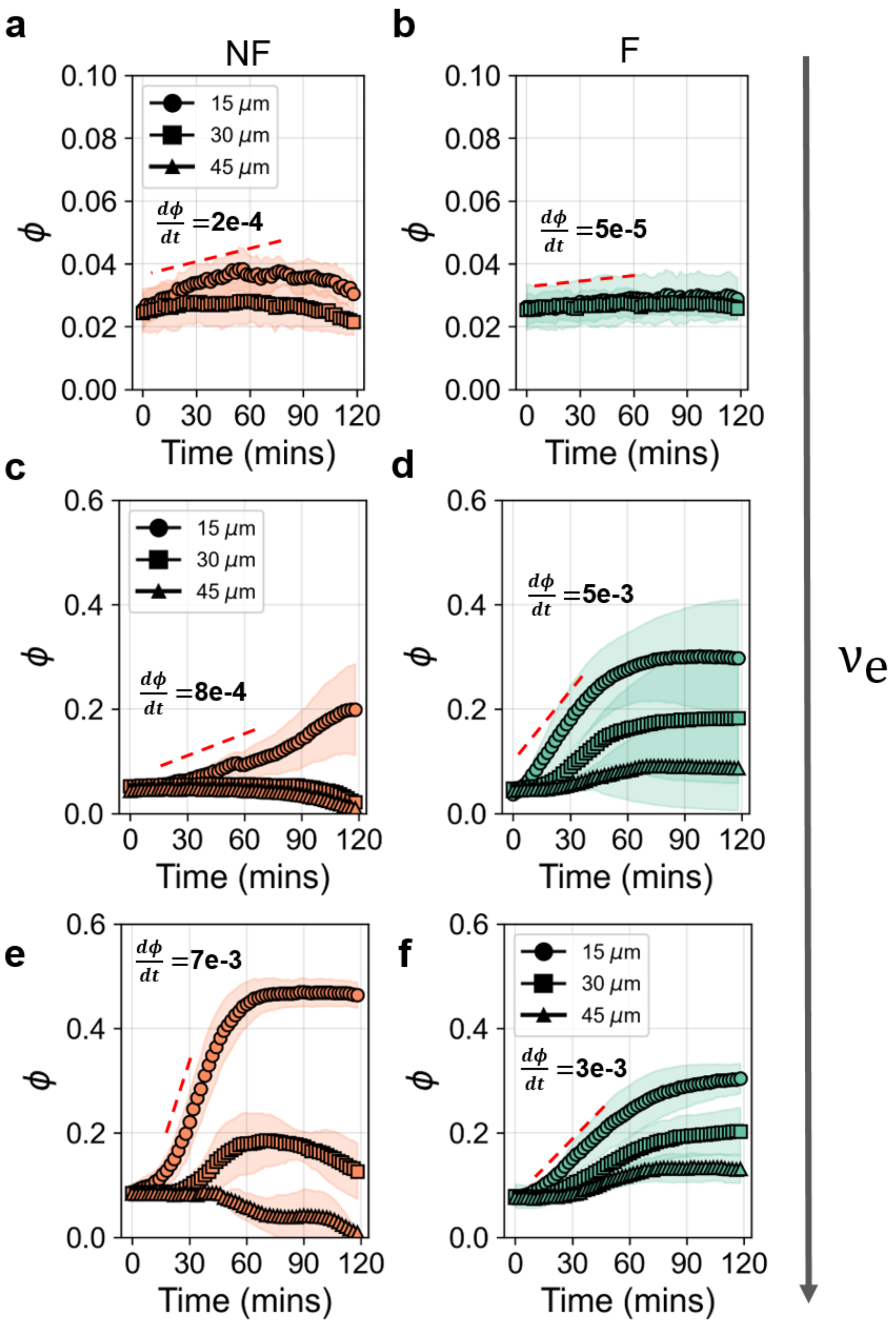


**Supplementary figure 5: $\phi$ vs time curves for each of the F and NF phenotypes and for each of the number densities tested.** Shown are respective $\phi$ vs time curves taken in increments of 15 $\mu$m from the bottom surface of the capillary and into the bulk volume of the capillary. The red lines correspond to the maximum gradient of the curves for the respective slices 15 $\mu$m above the bottom surface. The lowest number density displayed $\phi$ increases for the surface layer only, as no 3D network was formed. Shown are the mean and standard deviation from 3 biological replicates within which there were at least 2 different fields of view (441 $\mu$m $\times$ 441$\mu$m).

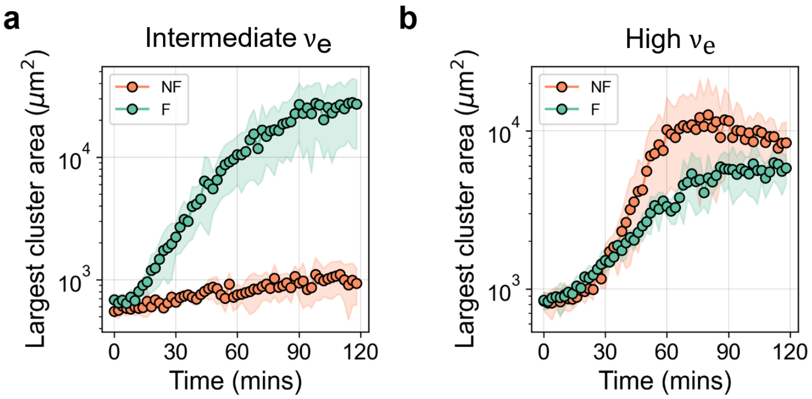


**Supplementary figure 6: Area of the largest bacterial clusters.** The size of the largest cluster for both the intermediate **(a)** and high number densities **(b)** for both the non-filamented (NF) and filamented (F) cells. The clusters were analyzed 30 $\mu$m above the bottom surface of the capillary from brightfield images. Shown are the mean and standard deviation from 3 biological replicates within which there were at least 2 different fields of view.

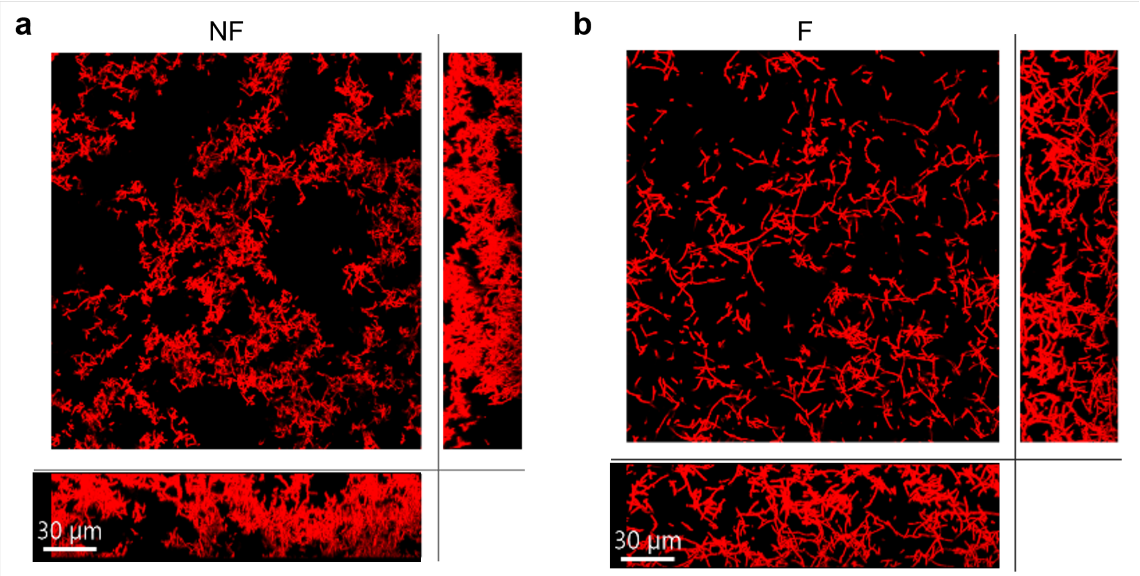


**Supplementary figure 7: Representative CLSM slices through the NF and F bacterial networks** $xz$ and $yz$ slices, with a 15$\mu$m projection depth depict the differing microstructure in the $z$ plane of the non-filamented **(a)** and filamented **(b)** bacterial networks. CLSM images were used to quantify branch lengths and the fractal dimensions

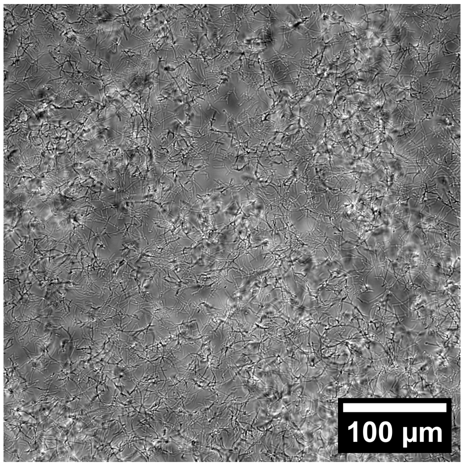


**Supplementary figure 8: Surface layer micrograph after delamination.** Representative micrograph of the filamented bacterial network surface layer of cells and "tufts" left on the capillary surface after a delamination event

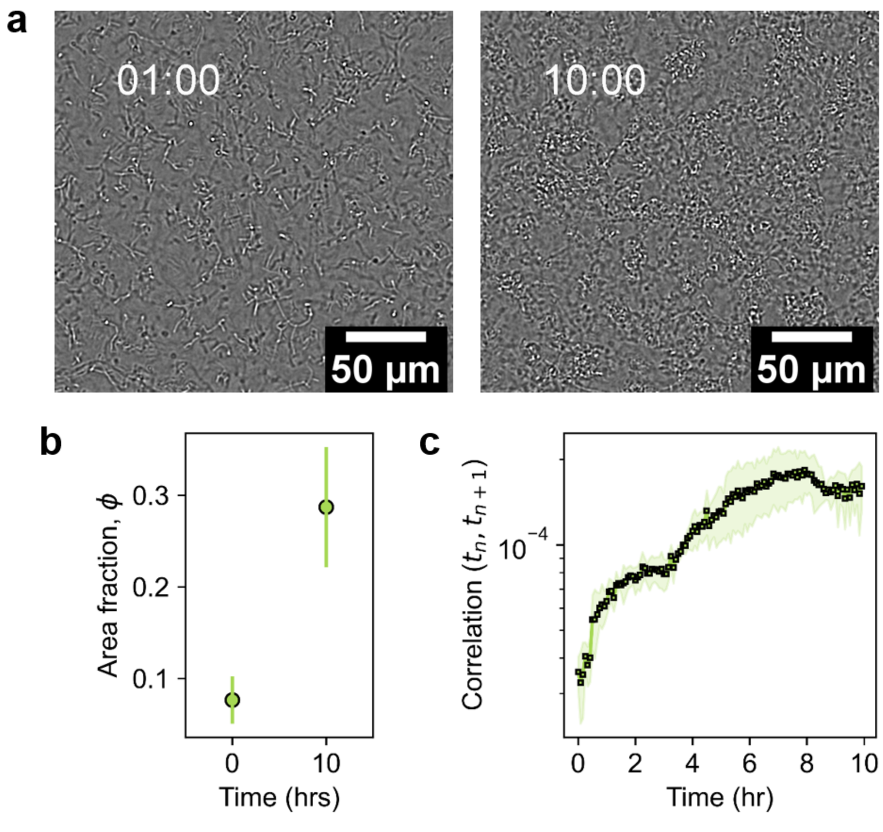


**Supplementary figure 9: Cluster-cluster aggregated networks density with growth and their dynamics are quench. (a)** Representative micrograph of the filamented bacterial network surface layer of cells after formation and after 10 hrs within a microfluidic channel **(b)** The area fraction of the networks increases with growth, plotted is the mean and s.d taken from two biological replicates. **(c)** Network fluctuations are quenched with growth and densification of the networks

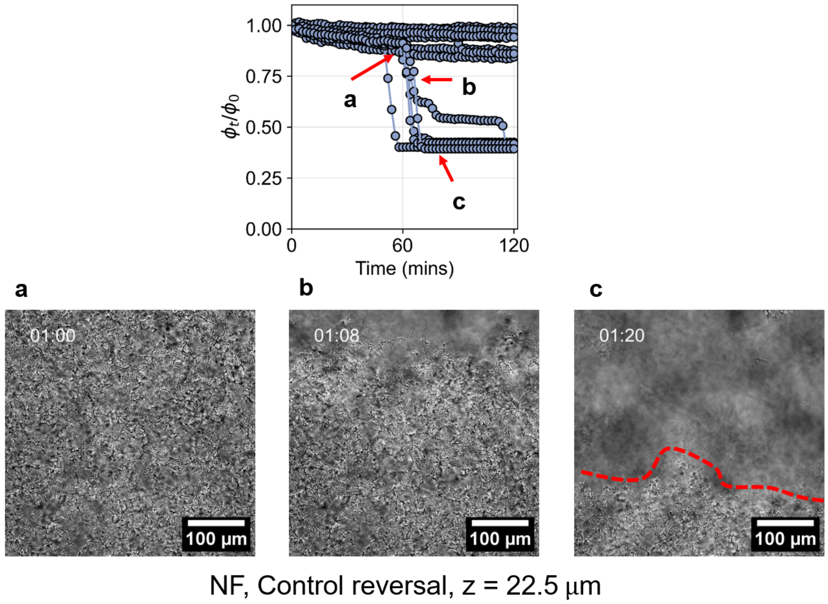


**Supplementary figure 10: Individual reversal curves for the NF control and representative fragmented yielding micrographs.** Shown are the normalised $\phi_t/\phi_0$ as a function of time for the NF bacterial networks. 5/15 of the networks display fragmented delamination behaviour, whilst the other 10/15 have type III behaviour and remain intact throughout the observation time. Fragmented yielding of an NF bacterial network, taken 22.5 $\mu$m beneath the top surface during a reversal experiment. **(a)** At t = 60 mins the network is intact, then a delamination event occurs at t = 68 mins **(b)**, finally ending with a fragmented structure where half of the network has delaminated and yielded from the network which remains adhered to the top surface of the capillary **(c)**.

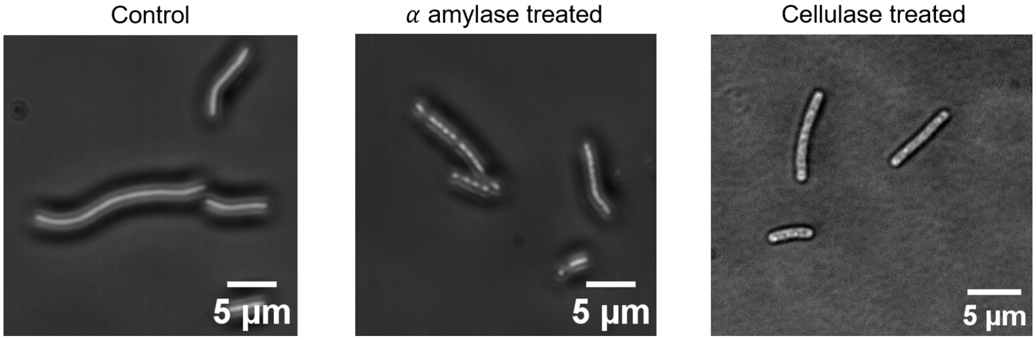


**Supplementary figure 11: Representative micrographs of *Comamonas denitrificans* F cells treated with glycoside hydrolase enzymes cellulase and $\alpha$ amylase.** The micrographs were taken in phase contrast mode using F cells after 2 hr enzyme exposure.

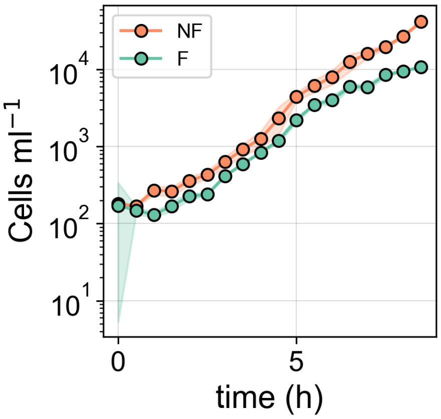


**Supplementary figure 12: Growth curves for *Comamonas denitrificans*.** Growth curves of the $O_2$ limited culture conditions, which induced the filamented phenotype (F) and of the $O_2$ replete culture conditions which induced the non-filamented (NF) phenotype. The growth rate was calculated from 5 hrs when the cultures exited the lag phase. Shown are the mean and standard deviation from 2 biological replicates.

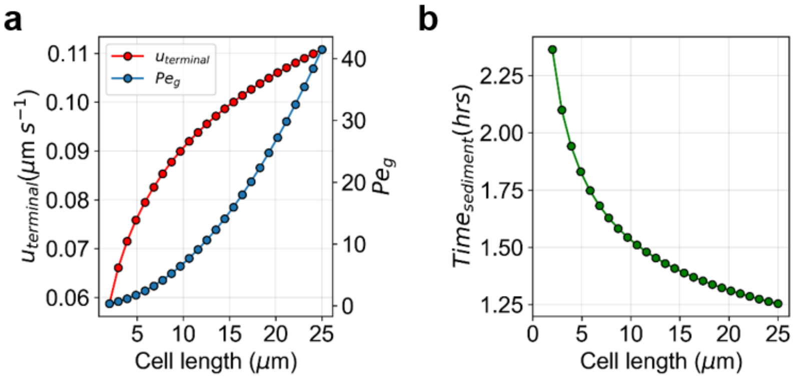


**Supplementary figure 13: Theoretical sedimentation calculations for *Comamonas denitrificans*. (a)** The theoretical terminal sinking velocity of *Comamonas denitrificans* as a function of cell length and the corresponding Peclet number, which indicates gravitational effects dominate over Brownian effects for the cell size range used in the experiments. **(b)** The theoretical time for a cell to sink the distance of the microscale capillary of 500 $\mu$m in height as a function of cell length, the cell density was assumed to be $\rho = 1.0825\ kg/m^3$.

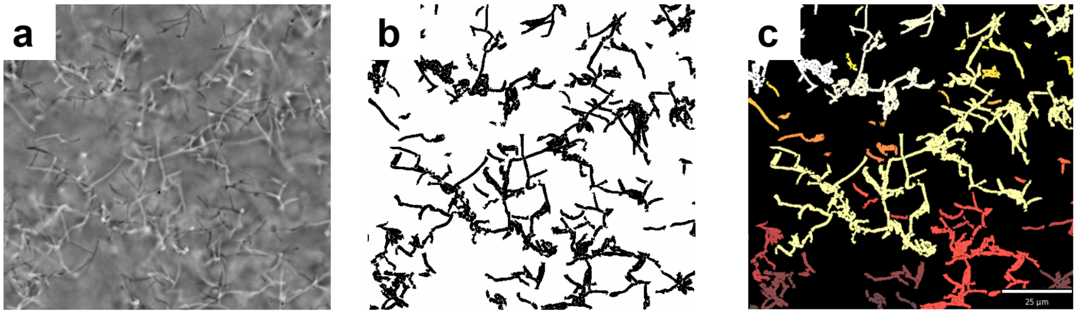


**Supplementary figure 14: Representative image processing steps.** (**a**) Raw images taken of a filamented (F) bacterial network, using brightfield microscopy. (**b**) The raw image is smoothed and then segmented using a subpixel curvilinear structure extraction algorithm (see Methods). (**c**) Processed images are then analysed using custom Python scripts to obtain network properties, the colors in this micrograph correspond to different clusters.

### Supplementary videos

**Supplementary video 1: Representative Yielding mode I of an untreated F bacterial network.** The video was acquired 30 $\mu$m below the surface of the imaging cell for 2 hours with a frame acquired every 2 mins.

**Supplementary video 2: Representative Yielding mode III of an untreated NF bacterial network**. The video was acquired 30 $\mu$m below the surface of the imaging cell for 2 hours with a frame acquired every 2 mins.

**Supplementary video 3: Representative non-yielding bacterial network.** Featured is an untreated NF bacterial network. The video was acquired 30 $\mu$m below the surface of the imaging cell for 2 hours with a frame acquired every 2 mins.

### Supplementary methods

#### 0.1 Growth curves

Growth curves were acquired by sampling from separate culture tubes every 30 mins for a period of 8 hours (Supplementary Fig. 10). In order to conserve the oxygen conditions tubes were sampled from once and then disposed of. 2 $\mu$L of culture was taken at each time point and added to 298 $\mu$L of 4% glutaraldehyde to fix the cells. At the end of the collection period, each of the wells was stained with 5 $\mu$M Syto 9 (Sigma Alderich, Switzerland). Cells were incubated for 15 mins before being transferred to the flow cytometer (cytoFLEX, Beckman Coulter, USA). Cells were counted in plate mode. Results from our live/dead assay indicated that staining with Syto 9 captured over 99% of the cell number from a stationary phase culture, indicating that staining with Syto 9 only would produce representative cell numbers of the population in the lag and exponential growth phase, which was captured during this experiment. The results were obtained using biological duplicates.

#### 0.2 Microfluidic network growth

To quantify the growth of the cluster-cluster assembled biofilms we used PDSM microfluidic channels (5 mm x 2 mm x 400$\mu$m, length x width x height). Channels were fabricated by pouring a 10:1 mixture of polymer and crosslinker onto an SU8 mask of the geometry. The PDMS was cured overnight at 80 °C. The microfluidic channels were cut, punched (with a 1 mm biopsy tool), and bonded to IPA-washed glass microscope slides using plasma activation. *C. denitrificans* was subcultured (100:1) from overnight cultures in 5ml TSB and grown until an optical density of 1 in a culture tube (13 ml). A syringe pump (Harvard Apparatus, USA) was used to control the flow into the channel. Briefly, the channels were purged with TSB media, to remove any air bubbles. Then the bacterial cultures were withdrawn into the channels from the channel outlet. The flow was stopped, and the system was left for 2 hours for networks to form. After two hours, a creeping flow was initiated (5 $\mu$L/hr), and the channels were imaged in two locations, up and downstream of one another ($\approx$ 500 $\mu$m separating distance) in the middle of the channel width. We performed time-lapse imaging in brightfield mode (Nikon Ti, Japan), with the microscope condenser 3/4 closed. Images were taken 15 $\mu$m above the bottom of the channel every 5 minutes for 10 hours. Two biological replicates were performed. To calculate the area fraction images were processed using ImageJ and Python. First, a difference in Gaussian filter was applied (using a radius of 1 and 2 pixels), then a variance filter was applied (radius 3). Finally, a simple threshold was applied to binarise the images. The area fraction was calculated as the number of pixels corresponding to 1 divided by the number of 0 pixels. The correlation coefficient was calculated from the non-binarised, variance-filtered images. Here a simple cross-correlation was applied with a time step of $\Delta$t = 5 mins, using the following equation: $\frac{(i_1 - \bar{x_{i1}}) * (i_2 - \bar{x_{i2}})}{std(i_1) * std(i_2)}$.

#### 0.3 Gravitational Peclet number

To non-dimensionalise the system the gravitational Peclet number is used, $Pe_g = V\Delta\rho gL/k_bT$, where $V$ is colloidal volume, $\Delta\rho$ is the density difference, $T$ is temperature and $k$ is Boltzmann's constant, which describes the ratios between gravitational force and thermal energy (Supplementary Fig. 11). The bacterial volume was approximated by assuming a sphero-cylinder geometry. A radius of 0.4 $\mu$m was taken from microscopy images and experimentally obtained cell length distributions were used. The density difference between water and the bacteria was measured using a percol gradient and found to be $\Delta\rho = 0.0835$ g/ml. The terminal velocity distribution of the suspensions was calculated using the following, assuming a fraction coefficient of $f_k = \frac{4\pi}{ln(\frac{2b}{a})-1/2}$. The volume of the cells was calculated as $V_{cell} = 4/3\pi r_{cell}^3 + \pi r_{cell}^2 l_{cell}$ and the mass as $m = \Delta\rho V_{cell}$.

#### 0.4 Cell characterisation: Live/Dead

The fraction of living and dead cells were measured using flow cytometry (CytoFLEX, Beckman Coulter, USA) . Briefly, cells were stained with Syto 9 (5$\mu$M)(Sigma Alderich, Switzerland) and Propidium iodide (20 $\mu$M) (Sigma Alderich, Switzerland) for 15 mins in PBS. Stained cells were diluted 1:1000 in PBS and then counted using flow cytometry.
